# Unraveling the Interactions between Human DPP4 Receptor, SARS-CoV-2 Variants, and MERS-CoV, converged for Pulmonary Disorders Integrating through Immunoinformatics and Molecular Dynamics

**DOI:** 10.1101/2023.03.06.531252

**Authors:** Arpan Narayan Roy, Aayatti Mallick Gupta, Deboshmita Banerjee, Jaydeb Chakraborty, Pongali B Raghavendra

## Abstract

Human coronaviruses like MERS CoV are known to utilize dipeptidyl peptidase 4 (DPP4), apart from angiotensin-converting enzyme 2(ACE2) as potential co-receptor for viral cell entry. DPP4, ubiquitous membrane-bound aminopeptidase is closely associated with elevation of disease severity in comorbidities. In SARS-CoV-2, there is inadequate evidence for combination of spike protein variants with DPP4, and underlying adversity in COVID19. To elucidate this mechanistic basis, we have investigated interaction of spike protein variants with DPP4 through molecular docking and simulation studies. The possible binding interactions between receptor binding domain (RBD) of different spike variants of SARS-CoV-2 and DPP4 have been compared with interactions observed in experimentally determined structure of complex of MERS-CoV with DPP4. Comparative binding affinity confers that Delta-CoV-2:DPP4 shows close proximity with MERS-CoV:DPP4, as depicted from accessible surface area, radius of gyration, number of hydrogen bonding and energy of interactions. Mutation in delta variant, L452R and T478K, directly participate in DPP4 interaction enhancing DPP4 binding. E484K in alpha and gamma variant of spike protein is also found to interact with DPP4. Hence, DPP4 interaction with spike protein gets more suitable due to mutation especially due to L452R, T478K and E484K. Furthermore, perturbation in the nearby residues Y495, Q474 and Y489 is evident due to L452R, T478K and E484K respectively. Virulent strains of spike protein are more susceptible to DPP4 interaction and are prone to be victimized in patients due to comorbidities. Our results will aid the rational optimization of DPP4 as a potential therapeutic target to manage COVID-19 disease severity.

## Introduction

DPP4 is a serine peptidase of highly conserved type II transmembrane glycoprotein, which comprises of a 6-residue N–terminal cytoplasmic tail, a 22 amino acid transmembrane, and extracellular domain (1). DPP4 is a diverse multifunctional protein, also known as T-cell activation antigen CD26 (2), or adenosine deaminase binding protein (ADBP) (3) The extracellular domain cleaves dipeptide after the second position from the N-terminus of peptides with proline or alanine, suggesting dipeptidyl peptidase activity (4). DPP4 is broadly distributed in the lung parenchyma, vascular endothelium, and fibroblasts of human bronchi, specifying that DPP4 may play vital role in modulating physiological and pathological functions in the lung (5).

Recently, DPP4 has obtained certain significance in the scenario of the severe acute respiratory syndrome coronavirus 2 (SARS CoV-2) infection, due to its ability as a cellular entry receptor or co-receptor for the virus. This is in support to the fact that DPP4 is well known receptor of MERS CoV (6). Such potential utilization of DPP4 as the binding target for MERS CoV offer us to predict the specific potential molecular interactions of SARS CoV-2 with DPP4. There has been an immense focus in the reported literature on the ability of both SARS CoV-2 and SARS-CoV to bind to angiotensin-converting enzyme II (ACE2) protein to enter the host cells (7, 8). ACE2 is considered as the primary receptor for spike protein of SARS CoV-2 to initiate infection (8). Since the outbreak, many studies have been published describing the distribution of ACE2 receptor in the different types of human cells, such as lung, liver, kidney, and colon (9), which indicates that SARS CoV-2 may infect different organs in the human body. However, the primary target cell for SARS CoV-2 entry is the lung alveolar type 2 (AT2) cells, which expresses reduced levels of ACE2 (9) conferring probable existence of co-membrane proteins facilitating host entry and infection. DPP4 interacts with several proteins that are important for viral processes and immune responses including ACE2, which implies a crosstalk between the two proteins that seeks further inspection.

The most prevalent comorbidities in SARS CoV-2 infected patients were hypertension and diabetes, followed by cardiovascular and respiratory diseases (10). Interestingly, DPP4 has a striking role in these disorders, especially on type 2 diabetes mellitus (T2DM). DPP4 play vital role in glucose homeostasis via proteolytic inactivation of peptides, such as glucagon-like peptide-1 (GLP-1), incretin hormones, and glucose-dependent insulinotropic polypeptide (GIP) (11). DPP4 concentration was demonstrated to be higher in individuals with obesity than in those with normal body weight (12). Barchetta et al. found higher circulating DPP4 activity in non-alcoholic fatty liver disease (NAFLD) patients than in non-NAFLD patients (13). DPP4 inhibitor decreases plasma apolipoprotein B and triglyceride levels in patients with T2DM, indicating the function of DPP4 in regulating lipid metabolism (14). DPP4 play critical role in pulmonary impairment. Chronic obstructive pulmonary disease (COPD) related impaired airflow imposes inflammatory response by DPP4. DPP4 turns on CXCL12 that may further activate proteases either directly or via chemokine regulation to exacerbate tissue degradation in COPD (15). DPP4 and a member of its gene family (DPP10) are also implicated in the pathophysiology of asthma (16,17). Interleukin-13 (IL-13), secreted by Th2 cells, has been shown to be related to airway inflammation and allergy in asthma (18). DPP4 expressed in pulmonary atrial smooth muscle cells mediate hypoxia induced pulmonary hypertension (19, 20). DPP4 has identified roles in other infections. In chronic hepatitis C virus (HCV), DPP4 generates an antagonist form of the chemokine CXCL10 (also known as IP-10) by amino-terminal truncation of the protein (21), such that the elevated plasma CXCL10 found in patients with chronic HCV can modulate immune responses by chemokine receptor antagonism (22). CXCR3 antagonism via truncated CXCL10 may also be an important regulatory mechanism occurring in tumours (23) and in sites of tuberculosis pathology (24). Exploring the role of such multi-faceted molecule in lung disease may yield vital and novel therapies.

Little attention is given to the cell-surface co-factors in facilitating the attachment and entry of SARS-CoV-2 currently. There are few computational studies based on the interaction of spike protein of SARS CoV-2 with DPP4 through molecular docking studies, however, effect of DPP4 interaction due to spike protein mutations have not yet explored. The varied emerging variants of SARS CoV-2 spike protein from (i) alpha, (ii) beta, (iii) delta, (iv) gamma and (v) omicron (the list of mutation in each variant is given in (Table S1) are considered in the current study to investigate the binding interaction with DPP4. Such finding may highlight the role of DPP4 in SARS CoV-2 variants and may elicit the effect of mutation on binding affinity by molecular docking and MD simulation approach. The results of our study will provide insight into the molecular mechanisms of viral entry and could have important implications for the development of new treatments for SARS-CoV-2. By comparing the predicted interactions between SARS-CoV-2 and DPP4 with those observed in MERS-CoV, our study seeks to clarify the potential role of DPP4 as a co-receptor for SARS-CoV2 and provide a basis for further investigation. We may analyse the effect of clustering of receptors like DPP4 and ACE2 both in association with SARS-CoV-2. From this we can predict about the variant associated role in Covid as well as in long Covid. SARS-CoV variants related DPP4 could have an influence on COPD, TB, Cardio-pulmonary diseases, and comorbidities. Further studies are needed to fully understand the complex interaction between DPP4 and MERS-CoV, as well as to develop strategies to target DPP4 in the prevention and treatment of MERS-associated SARS CoV-2 or comorbidities related diseases.

## Results

### Comparative analysis of spike variants and DPP4 interaction

To predict the specific binding potential of SARS CoV-2 with DPP4, molecular docking approach with HADDOCK has been followed. The overall orientation of the predicted binding interactions bear similarity to the predictions of Li et al., 2020 and Cameron et al., 2021 but do not reproduce their findings completely. A number of new, additional binding interactions are also noticed at the binding interface. The binding interactions of the docking pose most similar to the predictions by Li et al., 2020 (25) and Cameron et al., 2021 (26) (and indeed having the best interaction energy) is employed further for MD simulation study.

A total 500ns long MD simulations are carried out on DPP4 and spike protein variants from MERS CoV and SARS CoV-2 complexes. The final structure of each of the systems given by a snapshot is also shown as (Fig 1). The equilibrations of the simulated structures are assessed from the RMSD. RMSD is a quantitative measure of the variation of protein complex considering all the non-hydrogen atoms with respect to initial conformation of protein complex along the time of simulation, shown in supporting information, (Fig S1) We observe that all the systems equilibrate within 500ns of simulation time. The binding interactions between the RBD of SARS CoV-2 variants and DPP4 are compared to identify the most similar interactions to those of the MERS CoV:DPP4 complex. The changes in the binding state of different systems are manifested by several parameters.

**Figure 1:**
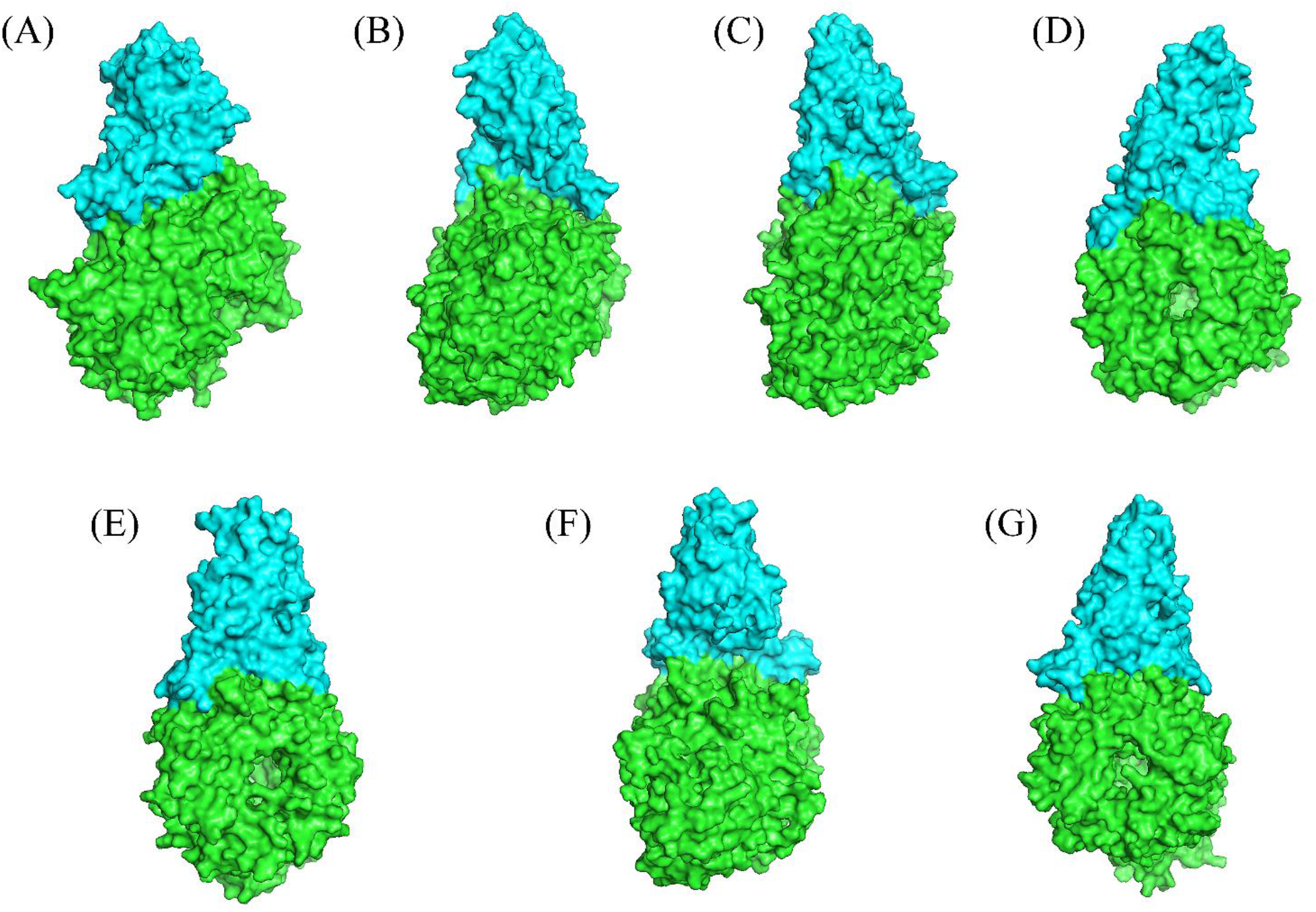
Snaphot (space-filled) of spike protein RBD (cyan) of (A) MERS (B) Wildtype (C) Alpha (D) Delta (E) Gamma (F) Omicron (G) Beta variant bound with DPP4 (green) from converged trajectory at 500ns.

### Radius of gyration

We characterize the comparative binding affinity of the complex through its radius of gyration (Rg). Rg of a complex measures its compactness. It is calculated as the average distance of the C-alpha atoms from their centre of mass. The Rg is computed for every ensemble and a histogram is generated for each system separately (Fig 2 (A, B)). Lower Rg in Delta SARS CoV2:DPP4 closely similar to MERS CoV:DPP4 complex, imparts more tight binding in them in comparison to rest of the ensemble.

**Figure 2:**
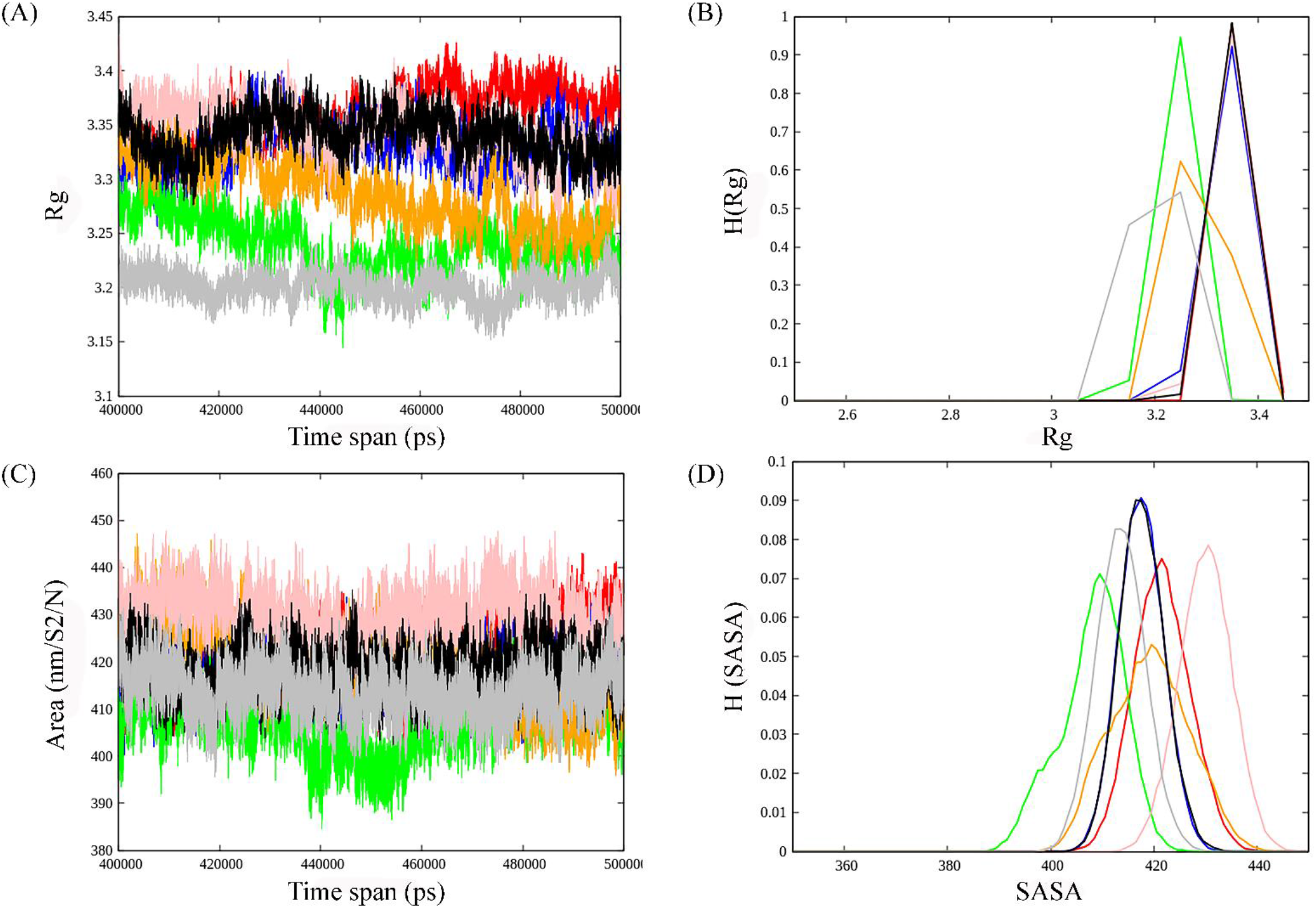
(A) Rg over time from converged trajectory, (B) Histogram distribution of Rg, (C) SASA over time from converged trajectory, (D) Histogram distribution of SASA. MERS-CoV:DPP4 is marked in grey, wild-type SARS CoV:DPP4 in black, alpha SARS CoV:DPP4 in red, beta SARS CoV:DPP4 in blue, delta SARS CoV:DPP4 in green, gamma SARS CoV:DPP4 in orange and omicron SARS CoV:DPP4 in pink.

### Accessible surface area

The contact surface at the binding interface depicted by solvent accessible surface area (SASA) is used as a descriptor to measure the relative DPP4 binding strength to various spike proteins. The less the surface area of a biomolecule in a complex that is accessible to a solvent elicits larger binding interface with higher buried surface area. Less accessible surface area for Delta SARS CoV-2:DPP4 and MERS CoV:DPP4 complex (Fig 2 (C)), confers increase in mean buried surface area at the binding interface indicating strong binding than that of the other interacting systems. The probability distribution of SASA, P(SASA) are shown in (Fig 2 (D)).

### Hydrogen bonds

The hydrogen bonds formed between DPP4 and spike protein variants of SARS CoV-2 and MARS CoV are calculated for all the 7 protein complexes. Such analysis gives the number of hydrogen bonds at a distance of less than 3.5Å between all possible donors D and acceptors A with a D-H-A angle of 180° to 30°. Overall, number of hydrogen bonds are quite high in MERS CoV:DPP4 and Delta SARS CoV-2:DPP4 complex (Fig 3 (A)), portraying stronger interaction in these systems than that of the rest. The distribution of the number of hydrogen bond, H(hb), is highest in Delta SARS CoV-2:DPP4 followed by MERS CoV:DPP4 (Fig 3 (B)).

**Figure 3:**
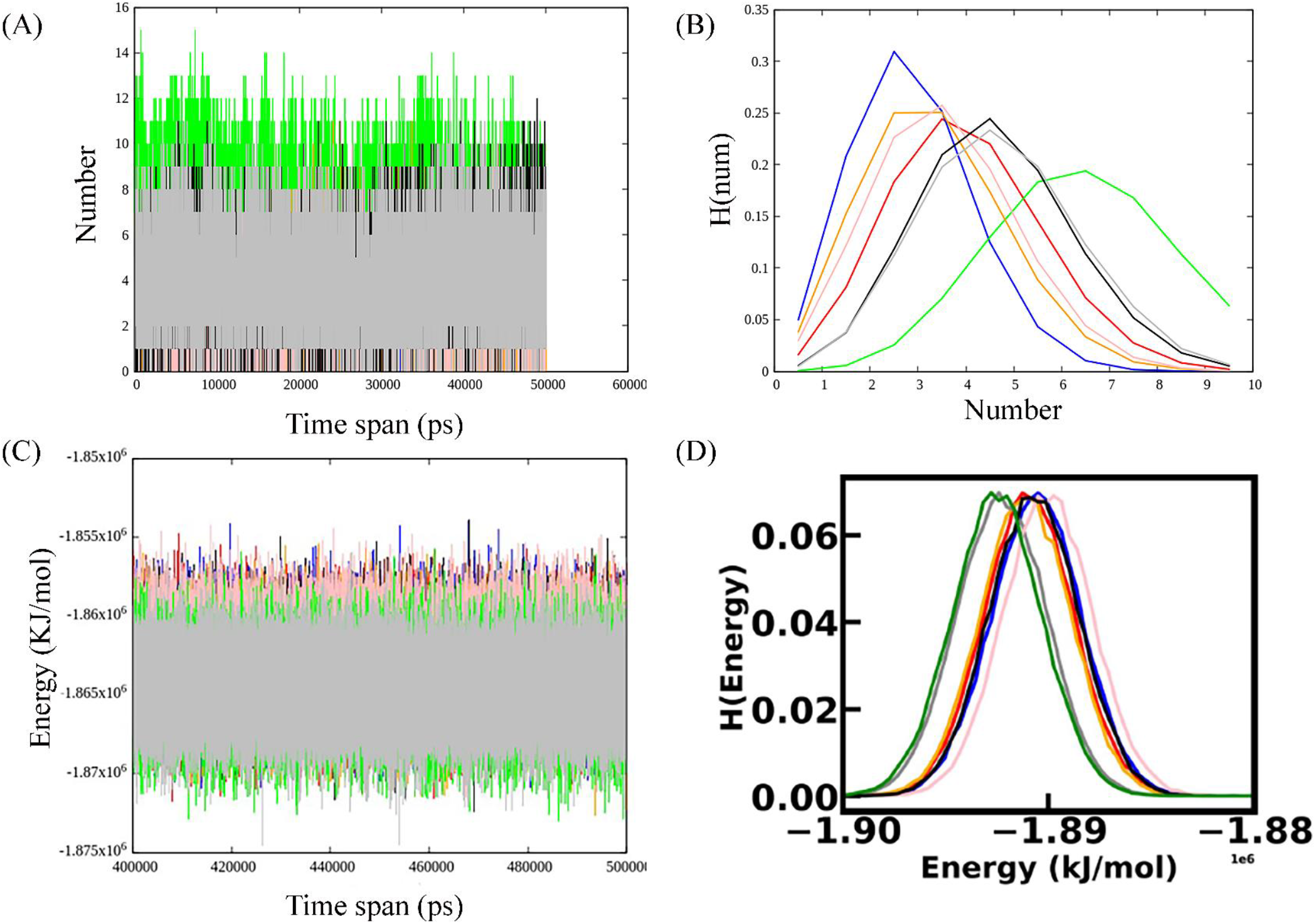
(A) Number of H-bond between chain A (DPP4) and chain B (spike) over time from converged trajectory, (B) Histogram distribution of H-bond number, (C) Interaction energy over time from converged trajectory, (D) Histogram distribution of Energy distribution. Color demarcations of the systems are same as Fig 2

### Interaction energy

We have also studied the total interaction energy between DPP4 and spike protein variants. A system becomes stabilized after attainment its minimum energy configuration. Thus, stabilization increases with a decrease in energy. MERS CoV:DPP4 and Delta SARS CoV-2:DPP4 is found to achieve least energy of interaction, hence, the most stable configuration is formed indicating stronger binding interaction (Fig 3 (C, D)).

### Flexibility

The intrinsic dynamics of the binding site have been analysed in our current study. Flexible and rigid regions of DPP4 and spike variants from each of the complexes are compared from RMSF and PCA calculations. RMSF considers the overall magnitude of the fluctuation of each Cα-atom, whereas PCA utilizes covariance matrix that includes information on the anisotropy of such fluctuations and their correlations between sites. PCA in MD simulation identifies only the essential motion eliminating other rotational and translational movements. Both RMSF and PCA analysis for DPP4 in all the complex is almost uniform (Fig 4 (A, B)). Overall fluctuation in DPP4 protein is not so profound. Perturbation in the spike protein variants due to RMSF is noticed in delta SARS CoV-2 and alpha SARS CoV-2. However, PCA of beta SARS CoV-2 is more fluctuating (Fig 4 (C, D)). This high degree of flexibility in the RBD region of delta variant is likely to be necessary for it to be able to interact with DPP4.

**Figure 4:**
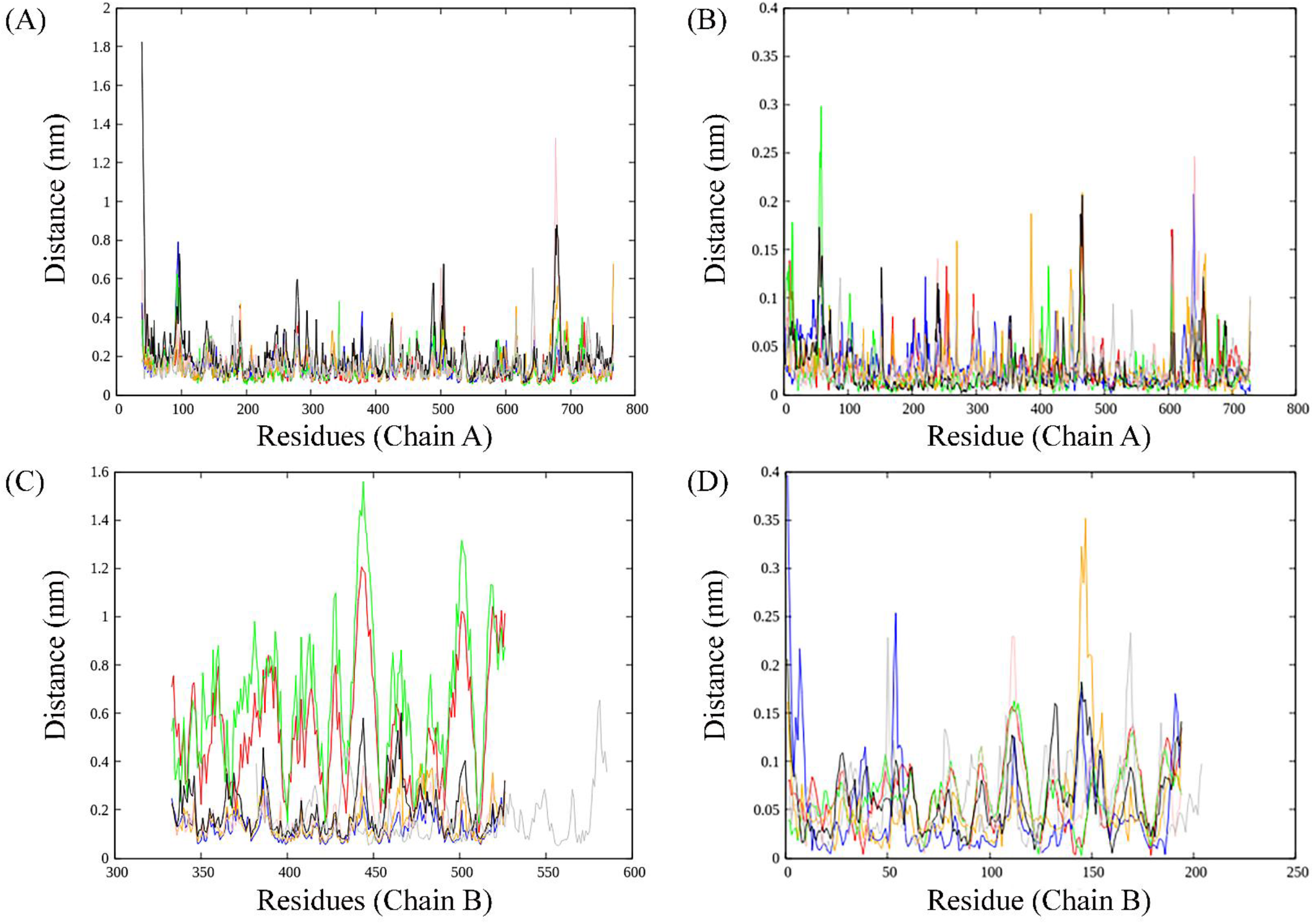
(A) RMSF and (B) PCA analysis for DPP4 (chain A) of the complexes. (C) RMSF and (D) PCA analysis for spike variants (chain B) of the complexes. Color demarcations of the systems are same as Fig 2

### Secondary structure

We have used DSSP to assign the secondary structural elements to the residues. The secondary structural elements play vital role in mediating interactions in the binding interface. Studies on the structural analysis and comparison of protein-protein interfaces depicts regular secondary structures (helices and strands) are the main components of the protein homodimer (obligate interface), whereas non-regular structures (turns, loops, etc.) frequently mediate interactions in the heterodimeric protein-protein interfaces (27). Secondary structural changes of DPP4 and spike protein for each of the ensemble from the converged trajectory have been studied vividly. We found secondary structure of DPP4 remains same in all the complex during various time frame of MD simulation (Fig 5). The spike protein, however, pertains substantial changes in secondary structure across the varied time span of simulation (Fig 6). In delta SARS CoV-2:DPP4, spike protein, non-regular secondary structural element, like bend is showing abundance in comparison to the rest. In comparison to MERS CoV:DPP4, the turn is found to decrease to certain extant in case of SARS CoV-2 variants. Spike protein of MERS CoV is less α-helical than that in the variants of SARS CoV-2. Prevalence of non-regular secondary structural elements in MERS CoV and delta SARS CoV-2 RBD region justifies it to be a potent binding partner to DPP4 to form the heterocomplex.

**Figure 5:**
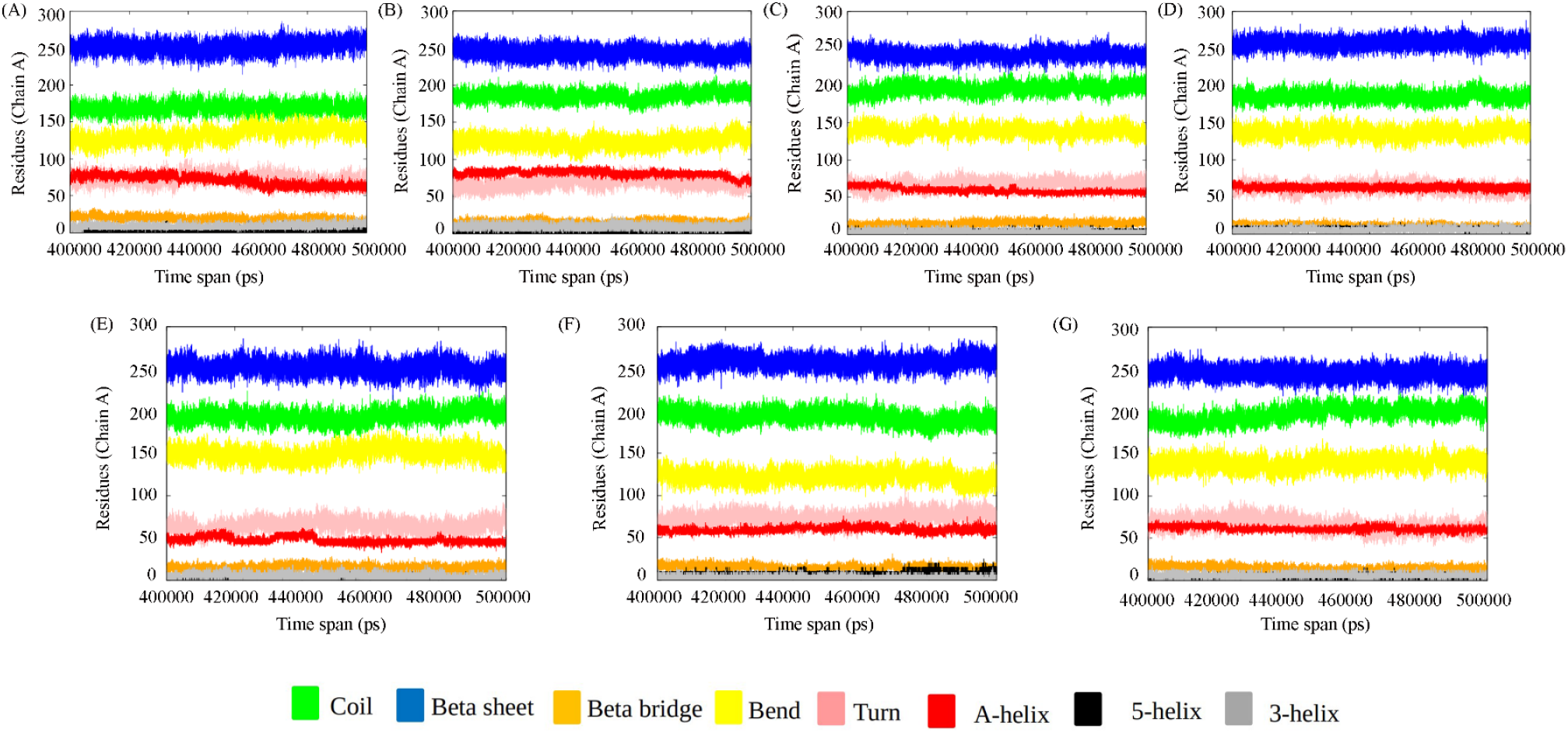
Secondary structure of DPP4 (protein chain A) of the complexes during various time frame of MD simulation. (A) MERS CoV:DPP4, (B) wild-type SARS CoV-2:DPP4, (C) Alpha SARS CoV-2:DPP4, (D) Beta SARS CoV-2:DPP4, (E) Delta SARS CoV-2:DPP4, (F) Gamma-SARS CoV-2:DPP4, (G) Omicron SARS CoV-2:DPP4.

**Figure 6:**
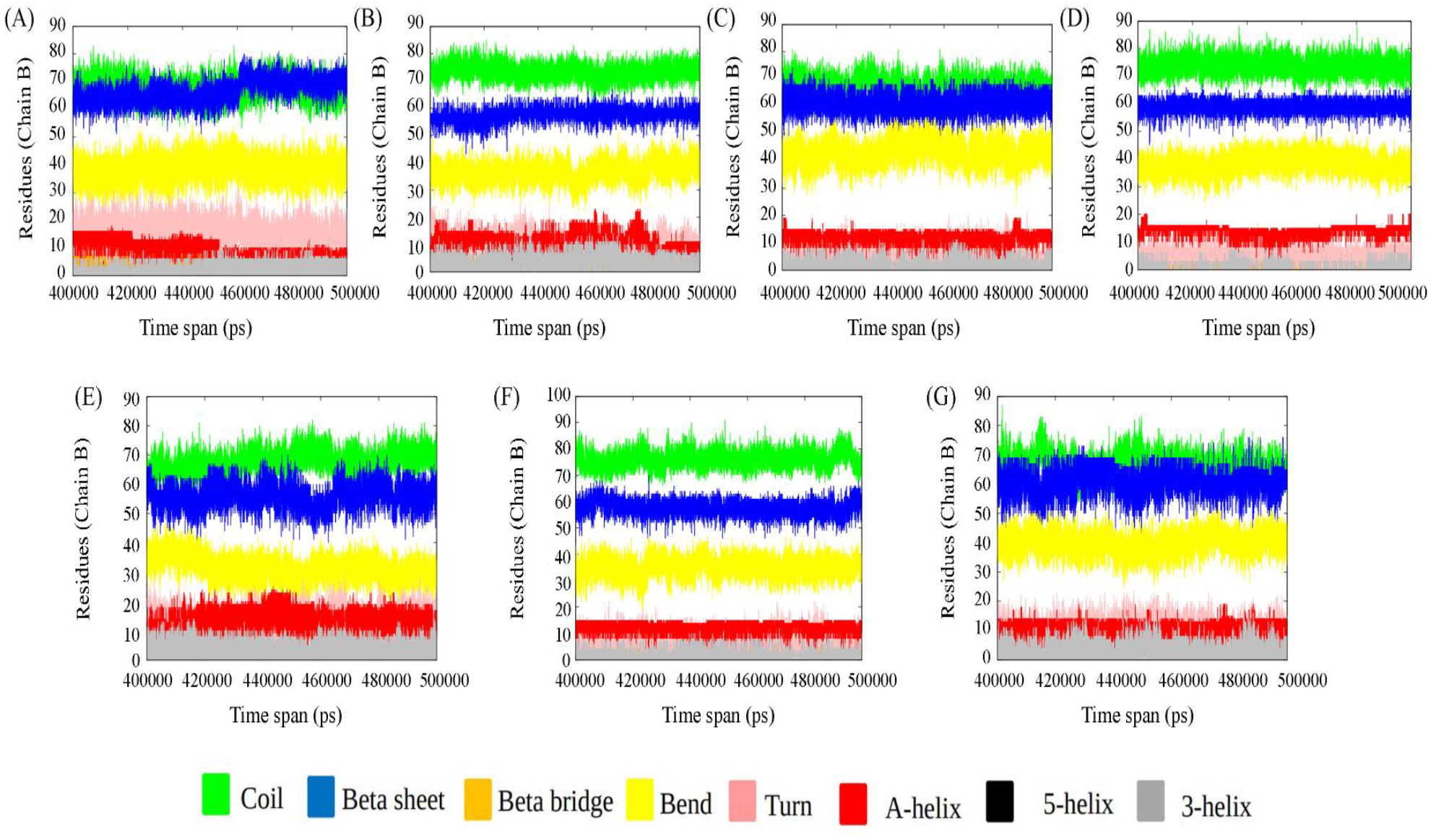
Secondary structure analysis of spike protein (protein chain B) (A) MERS CoV:DPP4, (B) wild-type SARS CoV-2:DPP4, (C) Alpha SARS CoV-2:DPP4, (D) Beta SARS CoV-2:DPP4, (E) Delta SARS CoV-2:DPP4, (F) Gamma-SARS CoV-2:DPP4, (G) Omicron SARS CoV-2:DPP4.

### Distance between centre of mass

We have also analysed the closeness between the two binding partner-DPP4 and spike from each of the ensemble. The binding interface have been studied by calculating the centre of mass of DPP4 and spike as a function of time. The protein-protein interface within 0.5 nm distance in each of the complex are tabulated (Table 1). The key residues forming the binding partner between MERS CoV and DPP4 as characterized from the x-ray diffraction crystal structures of their complex remain conserved till the end of the simulation (Table S2). Since SARS CoV-2 and MERS CoV are related virus, an important assumption in these studies is that RBD of SARS CoV-2 would likely bind to DPP4 as MERS CoV with similar conformation. Few studies have been conducted to demonstrate the SARS CoV-2 spike protein interaction with DPP4. The current study seeks to clarify the potential interaction of DPP4 with the different variants of SARS CoV-2 and compare it with MERS CoV through MD simulation study.

**Table 1:**
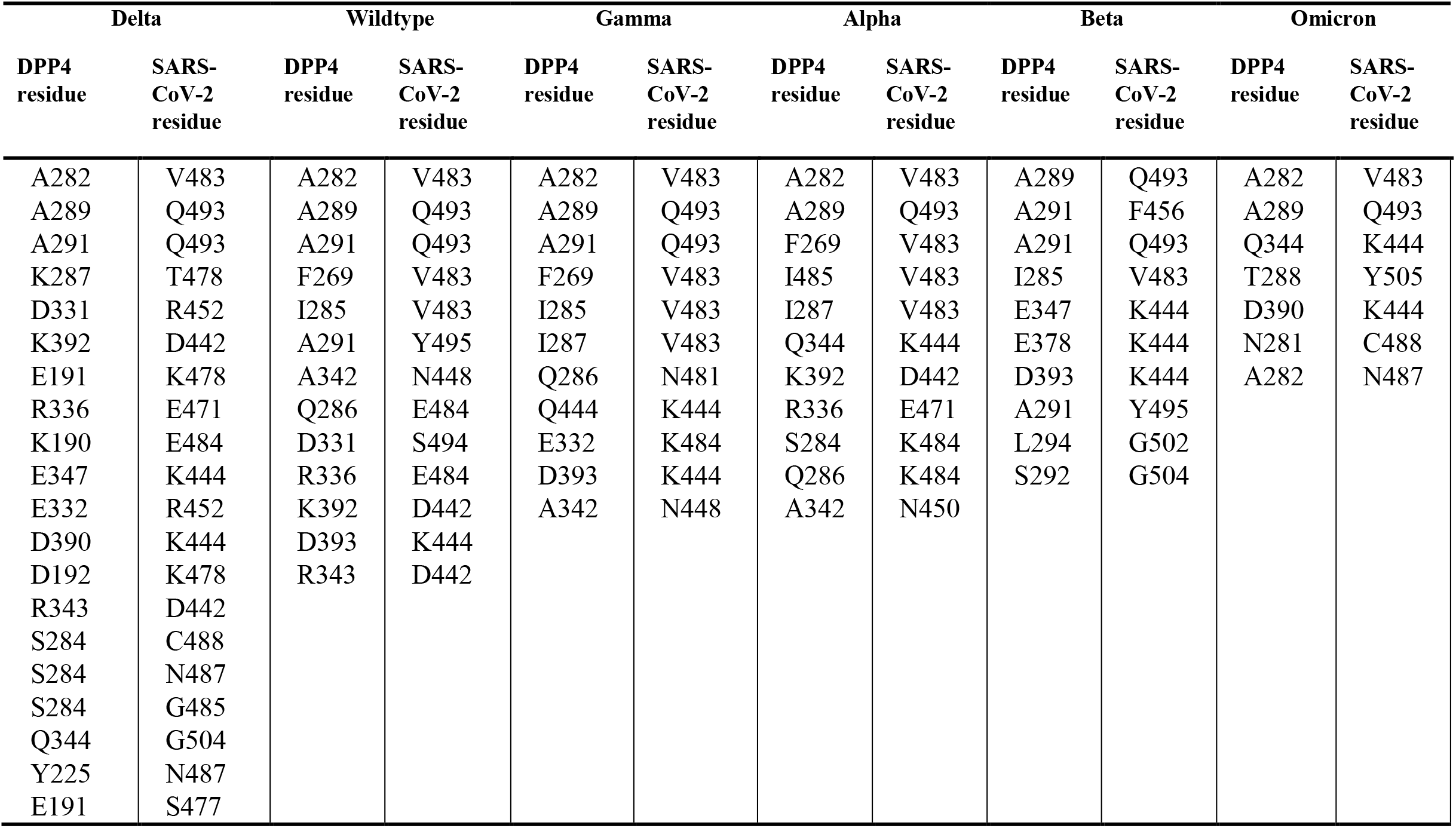
The binding interface at a distance of 0.5 nm between spike protein of SARS CoV-2 variants and DPP4

### Interface analysis

Interface analysis confers delta SARS CoV-2:DPP4 has more number of interaction than in other systems. The binding interface and the interacting partners with the mode of interactions are tabulated for each system (Table S3-S8). Our study has identified that A282 and A289 of DPP4 are found to interact with V483 and Q493 of spike variants respectively in all the cases. Thus, it forms the common interface. It is important to note that spike protein residues showing mutation in delta variant, L452R and T478K, directly participates in DPP4 interaction. Thus, mutation in delta variant enhances the binding. E484K in alpha and gamma variant of spike protein is also found to interact with DPP4. Thus, DPP4 interaction with spike protein gets more suitable due to L452R, T478K and E484K mutation. The more virulent strains of spike protein are more susceptible to DPP4 interaction and hence are prone to be victimized in patients due to comorbidities.

### Protein stability and flexibility

From the MD simulation study, we can observe that, spike protein residues showing mutation L452R and T478K in Delta variant, and E484K in Gamma and Alpha variant are found to directly participate in DPP4 interaction. Hence, we have chosen such interacting interface to analyse the effect of mutations on protein structures by calculating change in vibrational entropy (ΔΔS) and change in folding free energy (ΔΔG) as a thermodynamic state function. We have applied normal mode analysis (NMA) utilizing harmonic motions in a system, to provide insights into its dynamics and accessible conformations due to such mutations in the spike protein variants. The entropy and folding free energy changes are indicated in (Fig S2). In Delta variant L452R and T478K mutations are favoured by the attainment of molecular stability (0.692 Kcal/mol and 0.877 Kcal/mol respectively), and vibrational entropy changes (−0.154 kcal mol^-1^ K^-1^ and −0.728 kcal mol^-1^ K^-1^ respectively) leading to decrease in molecular flexibility. It is found that due to L452R mutation cconventional hydrogen bond between V350 and L452 breaks, and hydrophobic bond between L452 and L492 also breaks. In case of T478K, intra-molecular interaction remains same before and after mutation. On the other hand, E484K mutation for Gamma and Alpha variant elicits a destabilizing effect (−0.238 kcal/mol) on the protein and the vibrational entropy energy values (0.453 kcal mol^-1^ K^-1^) attributed towards increase of molecule flexibility. It appears that E484K mutation causes formation of one carbon hydrogen bond forms between K484 and C488.

### Perturbation residue scanning (PRS)

Additionally, to probe the effect of the interface mutations on the allosteric residue, PRS has been analysed. Perturbation effect can be analysed as a distance connecting the perturb residue to its nearby residues or on the interaction network strength. The PRS analysis of the most relevant interface mutations of the RBD of Delta (L452R, T478K), Gamma (E484K) and Alpha (E484K) has been done. In case of Delta variant, a perturbation analysis of R452 results in coupling distance of 3.3±0.2 Å and K478 results in coupling distance (d_c_) of 4.2±0.4 Å, and perturbation analysis of K484 (in case of Gamma and Alpha variant) reveals a d_c_ of 2.5±0.1 Å (Fig S3). If a mutation occurs at a residue with a high coupling distance, it is less likely to affect the stability of its interacting partner residues compared to a mutation at a residue with a low coupling distance. Therefore, in Delta variant the L452R mutation has maximum effect on the adjacent residues. We have also shown residue-wise Perturbation Residue Scanning profile (ΔQ) distance from the perturb site (Fig S4) as magnitude of perturbation is one of the factors in determining the effect of a mutation on adjacent residues. The coupling distance is uniquely sensitive to the environment of a residue in the protein to a distance of ~15 Å (28). We have also found out adjoining residues which are causing maximum perturbation with a cut-off distance 15Å due to mutation L452R, T478K and E484K (Fig S3). The maximum perturbation is observed in Y495 due to L452R mutation. Similarly, Q474 and Y489 shows significant perturbation due to the T478K and E484K respectively. Other factors, such as the location of the mutated residues within the protein structure and their interactions with other residues, can also affect the extent of perturbation and the resulting effect on adjacent residues.

## Discussion

This pandemic has become more dangerous as new SARS-CoV-2 variants have appeared in each new wave of infection. The primary functional receptor for SARS-CoV-2 is ACE2, DPP4 has also been suggested to be its potential co-receptor for viral cell entry (29, 30). In order to predict the specific potential interaction between DPP4 and various spike variants from SARS CoV-2 we have employed molecular docking and MD simulation studies. Notably, interaction between MERS-CoV spike protein and DPP4 is essential for viral infection and correlated with susceptibility to MERS-CoV infection, as well as with viral genome detection in the culture medium of infected cells (31). The binding interactions between MERS CoV and DPP4 have been characterized from their x-ray crystal structure of their complex. Such analysis of the binding interactions is very useful to compare from our MD simulation study. Interface analysis have shown that delta SARS CoV-2:DPP4 has a greater number of interaction than in other systems. Mutations can have varying effects on protein-protein interactions. Depending on the specific location and nature of the mutation, it can either strengthen or weaken the interaction between two proteins. In some cases, a mutation may create a new interaction site on one of the proteins or alter an existing interaction site, leading to stronger binding between the two proteins. This can result in enhanced biological activity or stability of the complex formed by the two proteins. The majority of these mutations have accumulated primarily in the Spike (S) glycoprotein. Since the Spike protein serves as the main infection target, focusing on S protein mutations increases transmission effectiveness and allows for resistance to neutralizing antibodies. Mutations on the RBD region of spike have been reported to increase affinity for ACE2 receptor. Current study elicits that spike protein residues showing mutation like L452R and T478K in Delta variant, and E484K in Gamma and Alpha variant directly engages in DPP4 interaction. Binding affinity for each of the system have been compared with MERS CoV and DPP4 interactions. Interpretation of DPP4-spike variant interactions from SARS Cov-2 using molecular simulation-based recognition can be an effective approach to determine binding affinity considering hydrogen bonding, energy of interaction, buried surface area, and distance between centers of masses in interaction between the two proteins. The overall interaction is well-built in the spike protein of delta variant with DPP4, as known from the interacting interface. Further, the effect of mutations has been predicted on the protein stability and flexibility through the vibrational entropy change (ΔΔS) and free energy change (ΔΔG) calculations. In Delta variant L452R and T478K both are having a stabilizing effect on DPP4 interactions. Loss of flexibility due to such mutations in Delta-SARS CoV-2:DPP4 concludes a better binding interaction between the two proteins. Overall, our study compared DPP4 interaction with MERS, Wildtype and rest all five variants of SARS-CoV-2 and identifies unique RBD residues crucial for the interaction with the DPP4 receptor. By comparing the predicted interactions between SARS-CoV-2 all variants and DPP4 with those observed in MERS-CoV, our study seeks to clarify the potential role of DPP4 as a co-receptor for SARS-CoV-2 and provide a basis for further investigation. We may analyze the effect of clustering of receptors like DPP4 and ACE2 both in association with SARS-CoV-2. From this we can predict about the variant associated role in Covid as well as in long Covid due to DPP4 binding. DPP4 circulates in the plasma, and has multidimensional role in immune-regulation, inflammation, oxidative stress, cell adhesion and apoptosis by targeting different substrates. DPP4 mediated comorbidities due to SARS CoV-2 infections among patients with diabetes, hypertension, cardiovascular diseases are more common. The more virulent strains of spike protein are more susceptible to DPP4 interaction and hence are prone to be victimized in patients due to the comorbidities.

## Materials and Methods /Experimental procedures

### Docking Studies

Docking between the spike protein of SARS CoV-2 in its native form (PDB id: 6M0J, chain B) and human DPP4 (PDB id: 4L72, chain A) has been done in HADDOCK (32). The docking protocol includes (i) rigid-body docking, (ii) semi-flexible refinement stage and (iii) final optimization in explicit solvent. In the initial rigid-body energy minimization stage typically 1000 complex conformations are generated. The best 200 structures are selected for optimization through semi-flexible simulated annealing in torsion angle, followed by final short restained molecular dynamic simulation in explicit solvent. Clustering is done using cut-off 7.5Å and a minimum cluster size to 4. Thus, 200 structures are clustered in 10 clusters. The clusters are analysed and ranked according to their average interaction energies (sum of E_elec_, E_vdw_, E_ACS_) and their average buried surface area. The resulting docked structures are sorted by minimum energy criteria and root mean square deviation (RMSD) clustering. The interface of the docked complex is analysed based on Cα distance (5Å) between binding partner residues.

### System Preparation

The crystal structure of MERS CoV spike protein complexed with human DPP4 (PDB id: 4L72) (6) is utilized for all-atom MD simulations study. For wt-spike:DPP4, we use the docked structure of wt-spike protein of SARS CoV-2 (PDB ID: 6M0J, chain B) (33), with that of DPP4 (PDB ID: 4L72, chain A). All the different Variants of Concern (VoC) of spike protein of SARS CoV-2 bound with DPP4 (likewise alpha-spike:DPP4, beta-spike:DPP4, delta-spike:DPP4, gamma-spike:DPP4 and omicron-spike:DPP4) are obtained by substituting the required amino acid alterations from the final structure of wt-spike:DPP4 at 500ns time span. The list of mutations for each variant are tabulated in details in (Table S1). Thus, we have considered total seven different systems for MD simulation studies.

### MD Simulation Studies

We perform molecular dynamics (MD) simulation of all the systems in explicit water. Spc216 (Simple Point Charge) water molecule (34), is used for solvation under a dodecahedron box with a minimum distance between the protein and the box 1.0Å. Counterions (Na^+^ and Cl^-^) are added to make the system electroneutral GROMACS 2018.6 program with GROMOS96 53a6 force field (35), has been applied at 300K and 1 atm pressure in an isothermal-isobaric ensemble under standard protocols using periodic boundary conditions and 2 femtosecond time-step. The longer-ranged particle-mesh Ewald method (36). The total number of particles is maintained to be same in all simulations (N=164301) to make the simulated ensembles equivalent. The simulation trajectories are calculated up to 500ns. The equilibrations of the simulated structures are assessed from the saturation of the root mean squared deviations (RMSD). Various analysis are carried out from the converged trajectory with tools in GROMACS to examine the system properties. The trajectories are being loaded for calculation and visualization using VMD (37).

### Protein Stability and Flexibility Analysis

The DYNAMUT tool (38), has been used to analyze the effect of all mutations, present on the RBD domain and outside the RBD (truncated S1 domain) of the spike protein, free energy changes (ΔΔG) and vibrational entropy difference (ΔΔS) were calculated. Protein flexibility and stability were examined on the complexes based on these calculations by utilizing the normal mode analysis (NMA) based elastic network contact model (ENCoM). ENCoM is an Elastic Network Contact Model that employs a potential energy function and includes a pairwise atom-type non-bonded interaction term to add an extra layer of information regarding the effect of the specific nature of amino acids on dynamics within the context of NMA. ENCoM tries to approximate ΔΔG through the calculations of the vibrational entropy (ΔS) of wild-type and mutant structures.

ΔS between two conformations (A, B) in terms of their respective sets of eigenvalues is given by: 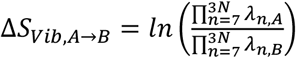

Where *λ_n,i_* represents the *n*th normal mode.

### Perturbation residue scanning (PRS)

PRS was performed using the pPerturb server (39, 40). It allows the mutation of one or more residues to alanine and generates a perturbation profile (ΔQ vs. C_alpha_-C_alpha_) distance from the perturb site. The ΔQ value is the magnitude of perturbation experienced by each residue and C_α_-C_α_ is the distance from the perturbed site. The perturbation effect can be analysed as a distance connecting the perturb residue to its nearby residues or on the interaction network strength. “Coupling distance” (d_c_) refers to a measure of the degree of coupling between two residues in a protein structure. This measurement is based on the interaction energy between residues and can provide insight into how changes in a protein’s structure can affect its function. The coupling distance can be used to predict the stability and functionality of a protein.

## Supporting information

The article contains supporting information.

## Acknowledgements

ANR gratefully acknowledge NIBMG, Kalyani facility for providing the technical support. AMG is thankful to the Technology Research Centre, S.N. Bose National Centre for Basic Sciences, Kolkata for the computational facilities.

## Author contributions

PBR, JC conceptualization; ANR, AMG, DB data curation, analysis; PBR, AMG, ANR methodology, software; AMG, ANR, DB acquisition, investigation resources; PBR funding acquisition, PBR, AMG writing – original draft; PBR, JC – review editing.

## Funding and additional information

We acknowledge NIBMG funding support, DB is thankful for CSIR - JRF fellowship funding, AMG is thankful to the Council for Scientific and Industrial Research for financial support through Research Associateship.

## Abbreviations

SARS: severe acute respiratory syndrome
MERS: middle East respiratory syndrome
ACE2: angiotensin-converting enzyme 2
DPP4: dipeptidyl-peptidase 4
MD: molecular dynamics
RBD: receptor binding domain
PDB: protein data bank
RMSD: root mean-square deviation
RMSF: root-mean-square fluctuation
SASA: solvent accessible surface area
PRS: perturbation residue scanning
PCA: principal component analysis
NMA: normal mode analysis
COPD: chronic obstructive pulmonary disease
T2DM: type 2 diabetes mellitus
GLP-1: glucagon-like peptide-1
GIP: glucose-dependent insulinotropic polypeptide
NAFLD: non-alcoholic fatty liver disease
HCV: hepatitis C virus
ADBP: adenosine deaminase binding protein

