## Supplementary Information for "Unraveling the Interactions between Human DPP4 Receptor, SARS-CoV-2 Variants, and MERS-CoV, converged for Pulmonary Disorders Integrating through Immunoinformatics and Molecular Dynamics"

### Figure legends

**Figure S1:** RMSD plot of (A) MERS CoV:DPP4, (B) wild-type SARS CoV-2:DPP4, (C) Alpha SARS CoV-2:DPP4, (D) Beta SARS CoV-2:DPP4, (E) Delta SARS CoV-2:DPP4, (F) Gamma-SARS CoV-2:DPP4, (G) Omicron SARS CoV-2:DPP4.

**Figure S2:** Visual representation of the interface mutations on dynamicity and plasticity of the RBD. RBD structure with the (A) L452R and (B) T478K mutation of Delta variant. RBD structure with the E484K mutation of (C) Gamma and Alpha variant. The RBD is represented as cartoon structure and mutations are shown as stick model. Red color in the cartoon structure indicates the flexibility in the protein and blue represents a rigidification of the structure. RBD, receptor binding domain

**Figure S3:** Distance dependence of the magnitude of perturbation on other residues ($\Delta Q$) from the perturbed site of (A) R452 & (B) K478 in Delta and (C) K484 in Gamma and Alpha

**Figure S4:** Residue-wise Perturbation Residue Scanning profile ($\Delta Q$) distance from the perturb site which is shown in dot in the graph shown for three interface mutations present on the RBD. (A) L452R (B) T478K (C) E484K (the residue that is perturbed is highlighted by a circle).

**Table S1**: List of mutations for each variant of SARS CoV-2. The crucial mutations are marked in bold.

| SARS CoV-2 Variants | Mutation |
|---|---|
| Alpha (B.1.1.7) | 69del, 70del, 144del, V70L, D178H, **E484K**, N501Y, P681H, A570D |
| Gamma (P.1) | N501Y, K417T, **E484K** |
| Beta (B.1.351) | N501Y, K417N, E484K, L18F, D80A, D215G, R246I, A701V |
| Delta (B.1.617.2) | **L452R**, **T478K**, G142D, 156del, 157del, R158G, D614G, P681R, D950N |
| Omicron (B.1.1.529) | K417N, N440K, G446S, S477N, T478K, E484A, Q493K, G496S, Q498R, N501Y, Y505H |

**Table S2:** The key residues forming the binding partner between spike protein of MERS CoV and DPP4

| DPP4 residue | MERS residue |
|---|---|
| K267 | D539 |
| R317 | D510 |
| R336 | Y499 |
| L294 | R542 |
| A289 | K502 |
| A291 | L506 |
| L294 | Y540 |
| L294 | V555 |

**Table S3:** The key residues forming the binding partner between spike protein of wildtype SARS-CoV-2 and DPP4

| DPP4 residue | SARS-CoV-2 Wildtype residue | Mode of interaction |
|---|---|---|
| A282 | V483 | Hydrophobic |
| A289 | Q493 | Hydrogen bond |
| A291 | Q493 | Hydrogen bond |
| F269 | V483 | Hydrophobic |
| I285 | V483 | Hydrophobic |
| A291 | Y495 | Hydrogen bond |
| A342 | N448 | Hydrogen bond |
| Q286 | E484 | Hydrogen bond |
| D331 | S494 | Hydrogen bond |
| R336 | E484 | Salt bridge |
| K392 | D442 | Salt bridge |
| D393 | K444 | Salt bridge |
| R343 | D442 | Salt bridge |

**Table S4:** The key residues forming the binding partner between spike protein of alpha SARS-CoV-2 and DPP4

| DPP4 residue | SARS-CoV-2 Alpha residue | Mode of interaction |
|---|---|---|
| A282 | V483 | Hydrophobic |
| A289 | Q493 | Hydrogen bond |
| F269 | V483 | Hydrogen bond |
| I485 | V483 | Hydrophobic |
| I287 | V483 | Hydrophobic |
| Q344 | K444 | Hydrophobic |
| K392 | D442 | Salt bridge |
| R336 | E471 | Salt bridge |
| S284 | K484 | Hydrogen bond |
| Q286 | K484 | Hydrogen bond |
| A342 | N450 | Hydrogen bond |

**Table S5:** The key residues forming the binding partner between spike protein of beta SARS CoV-2 and DPP4

| DPP4 residue | SARS-CoV-2 Beta residue | Mode of interaction |
|---|---|---|
| A289 | Q493 | Hydrogen bond |
| A291 | F456 | Hydrophobic |
| A291 | Q493 | Hydrophobic |
| I285 | V483 | Hydrophobic |
| E347 | K444 | Salt bridge |
| E378 | K444 | Salt bridge |
| D393 | K444 | Salt bridge |
| A291 | Y495 | Hydrogen bond |
| L294 | G502 | Hydrogen bond |
| S292 | G504 | Hydrogen bond |

**Table S6:** The key residues forming the binding partner between spike protein of delta SARS-CoV-2 and DPP4

| DPP4 residue | SARS-CoV-2 Delta residue | Mode of interaction |
|---|---|---|
| A282 | V483 | Hydrogen bond |
| A289 | Q493 | Hydrophobic |
| A291 | Q493 | Hydrophobic |
| K287 | T478 | Hydrophobic |
| D331 | R452 | Salt bridge |
| K392 | D442 | Salt bridge |
| E191 | K478 | Salt bridge |
| R336 | E471 | Salt bridge |
| K190 | E484 | Salt bridge |
| E347 | K444 | Salt bridge |
| E332 | R452 | Salt bridge |
| D390 | K444 | Salt bridge |
| D192 | K478 | Salt bridge |
| R343 | D442 | Salt bridge |
| S284 | C488 | Hydrogen bond |
| S284 | N487 | Hydrogen bond |
| S284 | G485 | Hydrogen bond |
| Q344 | G504 | Hydrogen bond |
| Y225 | N487 | Hydrogen bond |
| E191 | S477 | Hydrogen bond |

**Table S7:** The key residues forming the binding partner between spike protein of gamma SARS-CoV-2 and DPP4

| DPP4 residue | SARS-CoV-2 Gamma residue | Mode of interaction |
|---|---|---|
| A282 | V483 | Hydrophobic |
| A289 | Q493 | Hydrogen bond |
| A291 | Q493 | Hydrophobic |
| F269 | V483 | Hydrophobic |
| I285 | V483 | Hydrogen bond |
| I287 | V483 | Hydrophobic |
| Q286 | N481 | Hydrophobic |
| Q444 | K444 | Hydrophobic |
| E332 | K484 | Salt bridge |
| D393 | K444 | Salt bridge |
| A342 | N448 | Hydrogen bond |

**Table S8:** The key residues forming the binding partner between SARS-CoV-2 Omicron and DPP4

| DPP4 residue | SARS-CoV-2 Omicron residue | Mode of interaction |
|---|---|---|
| A282 | V483 | Hydrophobic |
| A289 | Q493 | Hydrophobic |
| Q344 | K444 | Hydrophobic |
| T288 | Y505 | Hydrophobic |
| D390 | K444 | Salt bridge |
| N281 | C488 | Hydrogen bond |
| A282 | N487 | Hydrogen bond |

### Figure S1

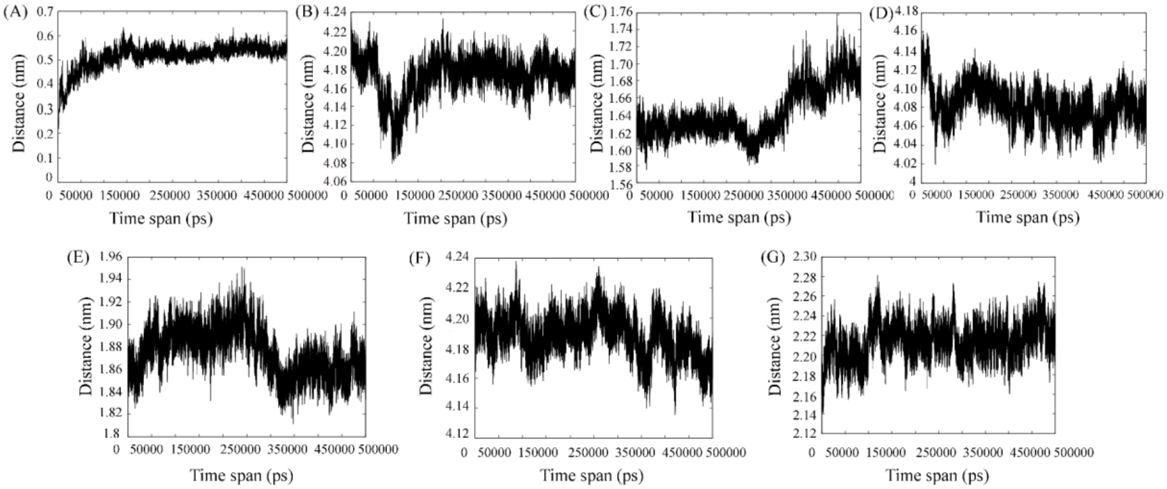

(A)
Distance (nm)
Time span (ps)
(B)
Distance (nm)
Time span (ps)
(C)
Distance (nm)
Time span (ps)
(D)
Distance (nm)
Time span (ps)
(E)
Distance (nm)
Time span (ps)
(F)
Distance (nm)
Time span (ps)
(G)
Distance (nm)
Time span (ps)

**Figure S2**

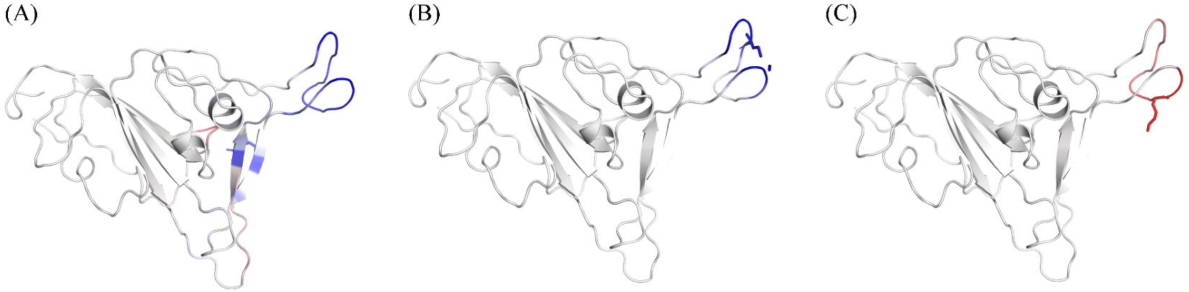

**Figure S3**

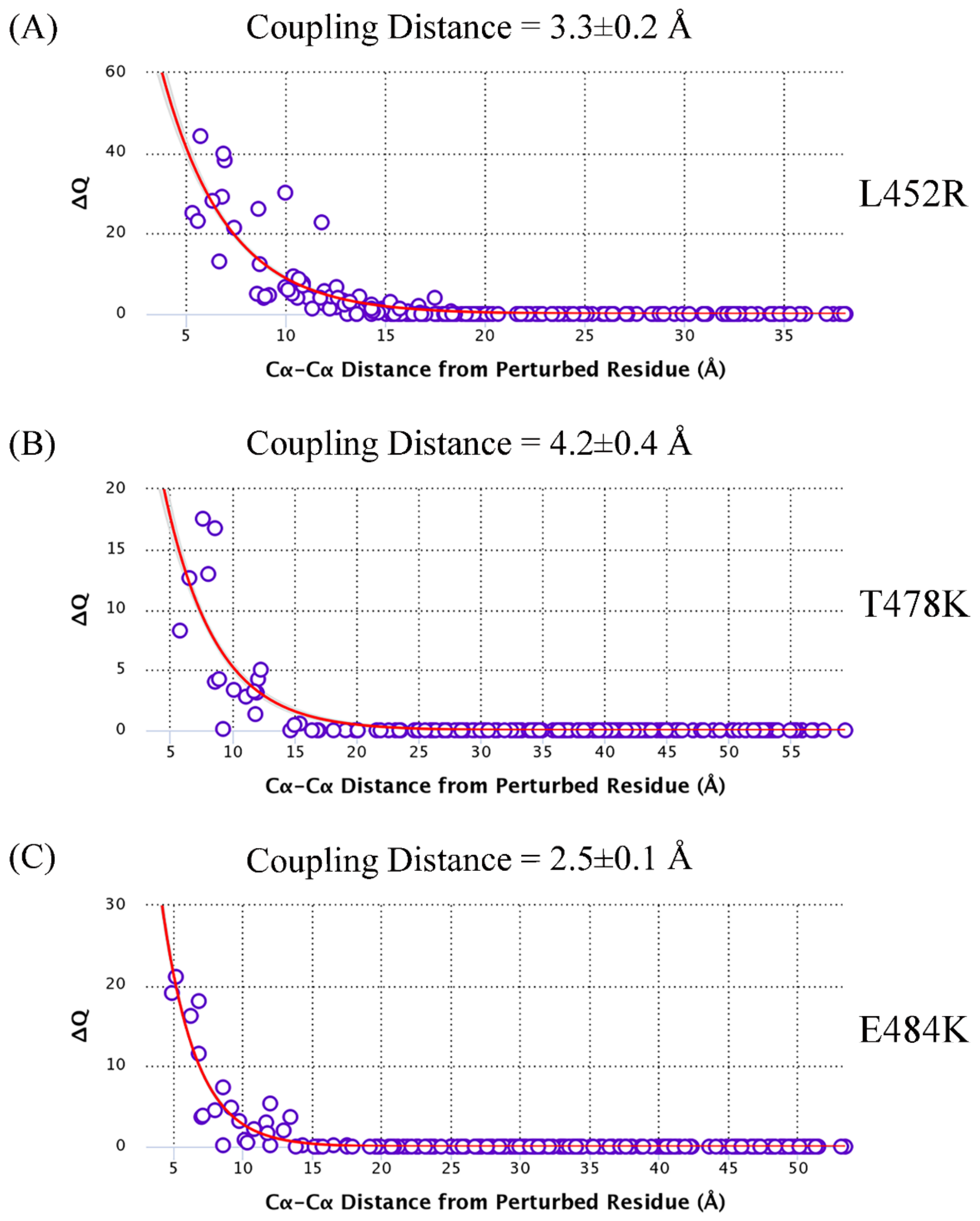

(A)
Coupling Distance = 3.3±0.2 Å
ΔQ
60
40
20
0
5
10
15
20
25
30
35
Cα-Cα Distance from Perturbed Residue (Å)
L452R
(B)
Coupling Distance = 4.2±0.4 Å
ΔQ
20
15
10
5
0
5
10
15
20
25
30
35
40
45
50
55
Cα-Cα Distance from Perturbed Residue (Å)
T478K
(C)
Coupling Distance = 2.5±0.1 Å
ΔQ
30
20
10
0
5
10
15
20
25
30
35
40
45
50
Cα-Cα Distance from Perturbed Residue (Å)
E484K

**Figure S4**

(A)

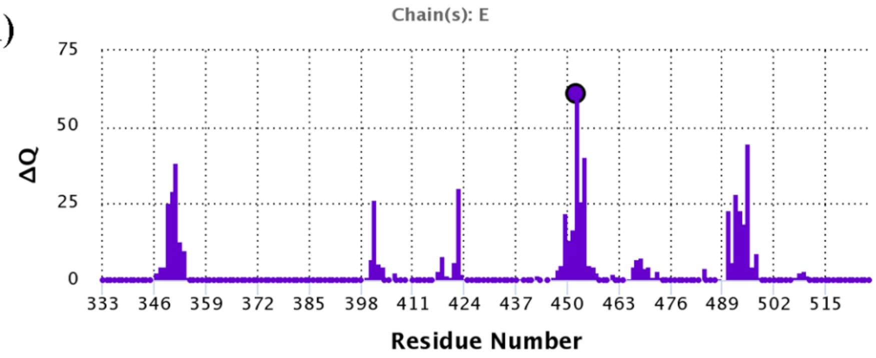


L452R

(B)

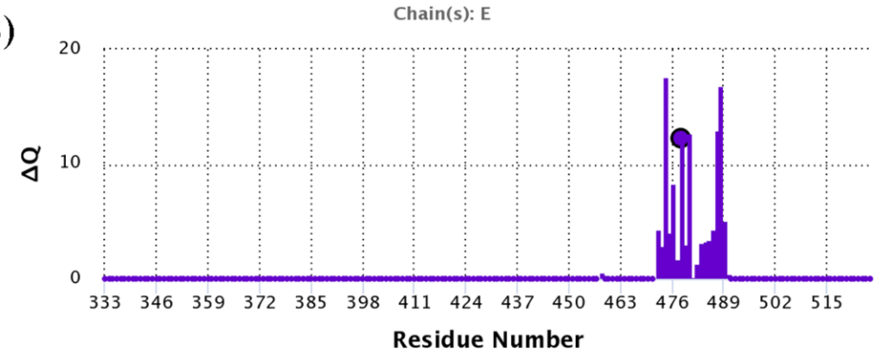


T478K

(C)

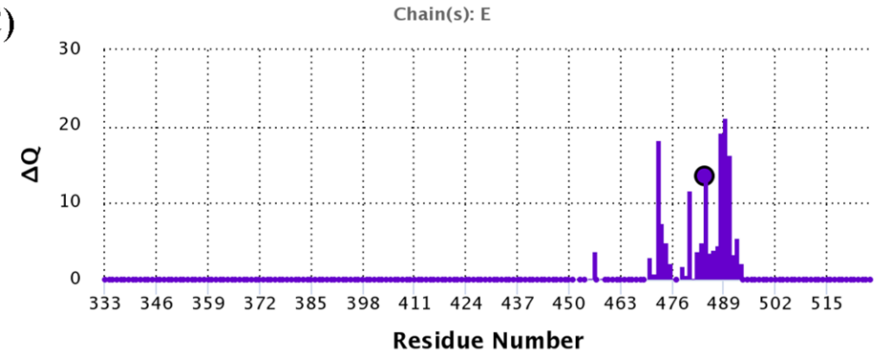


E484K
